# Label free 3D analysis of organelles in living cells by refractive index shows pre-mitotic organelle spinning in mammalian stem cells

**DOI:** 10.1101/407239

**Authors:** Patrick A. Sandoz, Christopher Tremblay, Sebastien Equis, Sorin Pop, Lisa Pollaro, Yann Cotte, F. Gisou van der Goot, Mathieu Frechin

## Abstract

Holo-tomographic microscopy (HTM) is a label-free non-phototoxic microscopy method reporting the fine changes of a cell’s refractive indexes (RI) in 3D. By combining HTM with epifluorescence, we demonstrate that cellular organelles such as Lipid droplets and mitochondria show a specific RI signature that distinguishes them with high resolution and contrast. We further show that HTM allows to follow in unprecedented ways the dynamics of mitochondria, lipid droplets as well as that of endocytic structures in live cells over long period of time, which led us to observe to our knowledge for the first time a global organelle spinning occurring before mitosis.

## Introduction

Because of the transparent nature of a cell, microscopy techniques either use fluorescent markers or transform optical properties of the sample into an observable contrast (*e.g.* phase contrast, DIC). Each of these techniques comes with limitations. Photobleaching, phototoxicity, interference of markers or ectopically expressed engineered proteins are major concerns. Classical label-free imaging techniques, while less perturbing, provide images with low information content due to poor contrast and resolution.

In this context, holo-tomographic microscopy (HTM)(1) is of great interest, as it provides label-free, high-content images without phototoxicity. HTM is a quantitative phase microscopy method(2–5) where the object’s complex wave field is encoded into a hologram. To do so, a reference beam and a beam that has interacted with the object are brought to interference on a CMOS camera(1–4). Moreover, HTM combines this holographic approach with rotational scanning of the specimen(5) (Fig 1A). The direct synthesis of thus gained series of scattering spectra offers a fast 3D reconstruction of even live(6) sample’s refractive index (RI) distribution at a resolution below the diffractive limit of light defined by the Rayleigh criterion(7). While RI distributions start to be used in life sciences studies(8–11) and specific refractive index signatures for cell structures(12,13) have been partially analyzed, current HTM systems typically suffer from coherent noise created by scattered light from the rotational scanning mechanism such as digital micro-mirror devices. This coherent noise impedes high spatial resolution and results in reduced RI sensitivity. Thanks to a completely scattering-free optical setup, the here presented HTM setup effectively overcomes this limitation and permits to our knowledge, the first systematic and precise characterization of cellular organelles by refractive index in space and time. We then use such capabilities to investigate the co-dynamics of cellular organelles over the cell cycle of mouse embryonic stem cells (mESCs) and shine light on a new phenomenon of organellar spinning implicated in cellular reorganization before mitosis.

**Fig 1.**
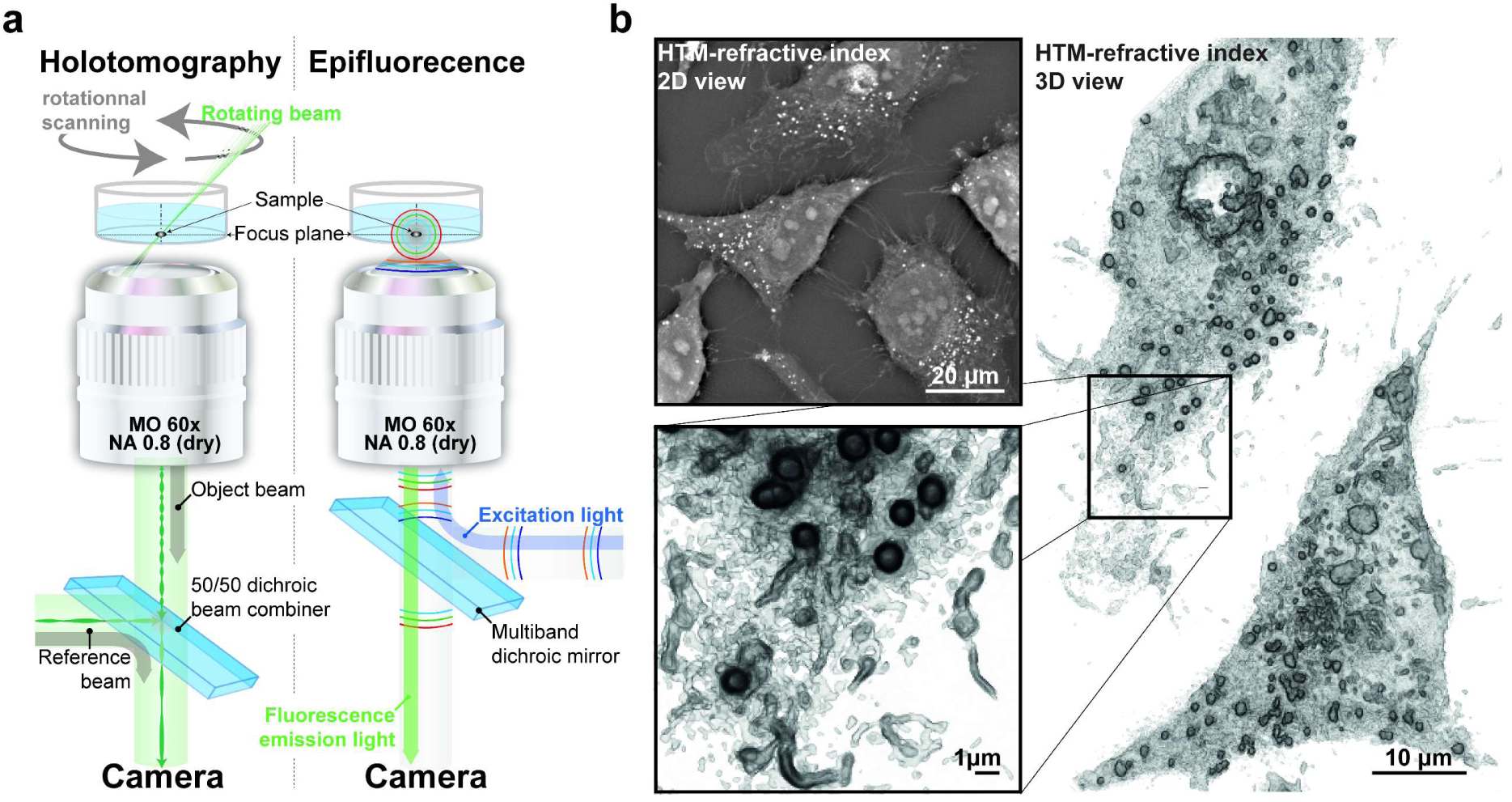
Holotomographic microscopy of subcellular structures. **a**, Scheme of the holotomographic microscopy (HTM) setup coupled to epifluorescence used in this study. **b**, 2D and 3D images of unperturbed HeLa cells taken with HTM.

## Results & Discussion

Using a novel device equipped with simultaneous holotomography and epifluorescence imaging (Fig 1A), we compared fluorescence imaging of subcellular markers with RI to determine whether specific organelles have a recognizable RI signature that would allow label-free analysis under control conditions and upon perturbation.

To compare HTM with other label-free methods, fixed HeLa cells were first imaged with bright-field, phase contrast DIC and HTM techniques (S1 Fig). Dehydrating fixation methods affected organelles and thereby their RI, but paraformaldehyde allowed their preservation (S2 Fig). The plasma membrane, the nucleus and lipid droplets (LDs) could be distinguished unambiguously by all technologies but were perfectly observable in 2D and 3D images provided by HTM due to their very specific RI signature (Fig 1B). LD were also visible by HTM in U2OS, Fibroblasts, RPE1, MEF, Hepa1.6, HAP1 and 3T3 cells differentiated in adipocytes (S3 Fig). In addition to LDs, HTM could clearly discriminates (in 2D and 3D) other intra-cellular structures that could not be distinguished by the other methods (Fig 1B, S1 Fig).

We next used the HTM/epifluorescence setup to compare the RI sections and the corresponding fluorescent signals of the cells under investigation. We first analyzed the largest organelle in the cell, namely the endoplasmic reticulum (ER) using KDEL-GFP as a fluorescent marker. Analysis of the corresponding RI map (S4A Fig), and the quantitative evaluation of signal correlations (S2B Fig) indicated that this organelle does not lead to a RI signal in control cells. Similarly, using NAGTI-GFP as a fluorescent marker, we found that the Golgi apparatus is not detected by HTM. The fact that the ER and the Golgi are “silent” in terms of RI mapping is advantageous given that the ER is present throughout the cell and its detection would have hampered the proper analysis of any other organelle. In contrast, labeling of cells with the mitochondrial marker Mito-YFP (Fig 2A) or lipid marker Bodipy (Fig 2B) revealed that mitochondria and lipid droplets provide a clearly distinguishable RI signal in 2D and 3D.

**Fig 2.**
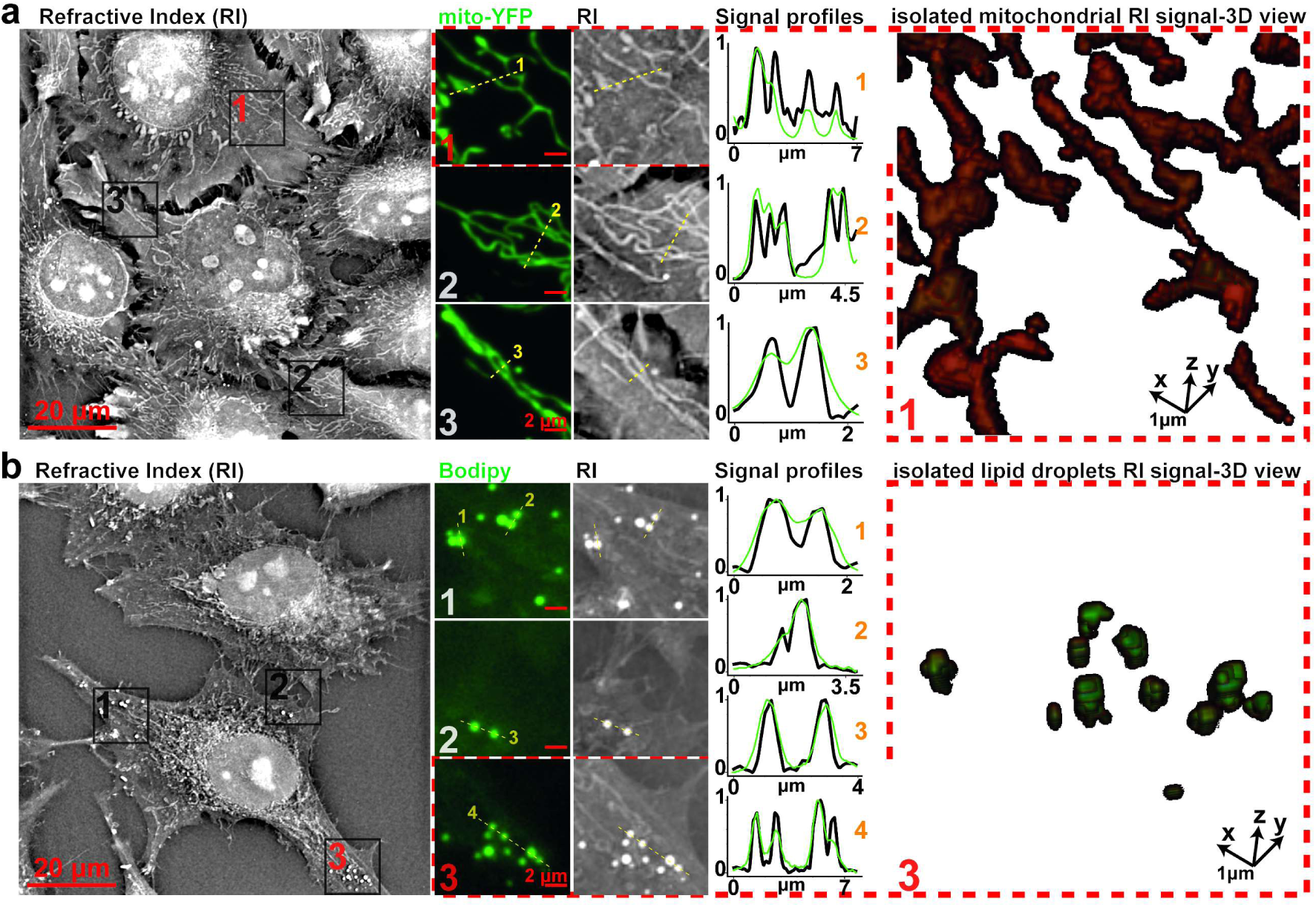
Comparison of specific fluorescent signals and refractive index map. Visual and profile comparison as well as 3D view of **a**, HeLa cells refractive index (RI) map to mitochondria-specific fluorescent signal (mito-YFP) or **b**, to a lipid droplet-specific fluorescent signal (Bodipy) shows specificity and gain in resolving power. RI and fluorescent signal profiles have been normalized between 0 (background) and 1 (profile maximum).

This visual conclusion was confirmed using Pearson coefficient and Kolmogorov-Smirnov tests. It was also confirmed by the comparison between fluorescent signals with independent human expert labelling of RI maps that shows over than 90% of overlap (S5 Fig), altogether validating a specific correlation of the RI and fluorescent signals. Interestingly, while in epifluorescence (or in a normal microscope using the same optics), such subcellular structures are being diffraction limited to maximum of resolution of 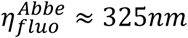, implicating that we cannot distinguish them properly, our HTM’s theoretical resolution limit(6) allows for 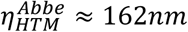. This property allows to finely resolve mitochondria or similar structure with a high RI precision(6) of 2.71E^-04^. Accordingly, the signal profiles in Fig 2A and 2B show a striking improvement in resolving subcellular structures using our RI measurement over the conventional fluorescence signal. Thus, the resolution of mitochondrial tubules down to 186 nm by the used HTM setup (S6 Fig) demonstrates a sub-diffraction imaging regime with respect to 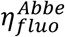. To further confirm the ability of HTM to analyze mitochondria, we perturbed mitochondrial fission by overexpressing Fis1(14). While cells expressing a mitochondria-targeted dsRed protein (S7A Fig) exhibit regular elongated and thin mitochondria, cells transfected with Fis1 presented perinuclear collapsed and round shaped structures (S7B Fig).

A strong asset of HTM is the extremely low amount of energy that is transferred to the sample during acquisition (0.01 nW/µm^2^ in presented setup), which guaranties the absence of phototoxicity. Therefore, HTM offers unique possibilities to observe truly unperturbed processes in live cells. This can be seen in our time-lapse acquisitions of mouse embryonic stem cells at a frequency of 4 image.min^-1^ that reveal that individual mitochondrial fission and fusion events can be monitored (Fig 3A and S1-3 movie).

**Fig 3.**
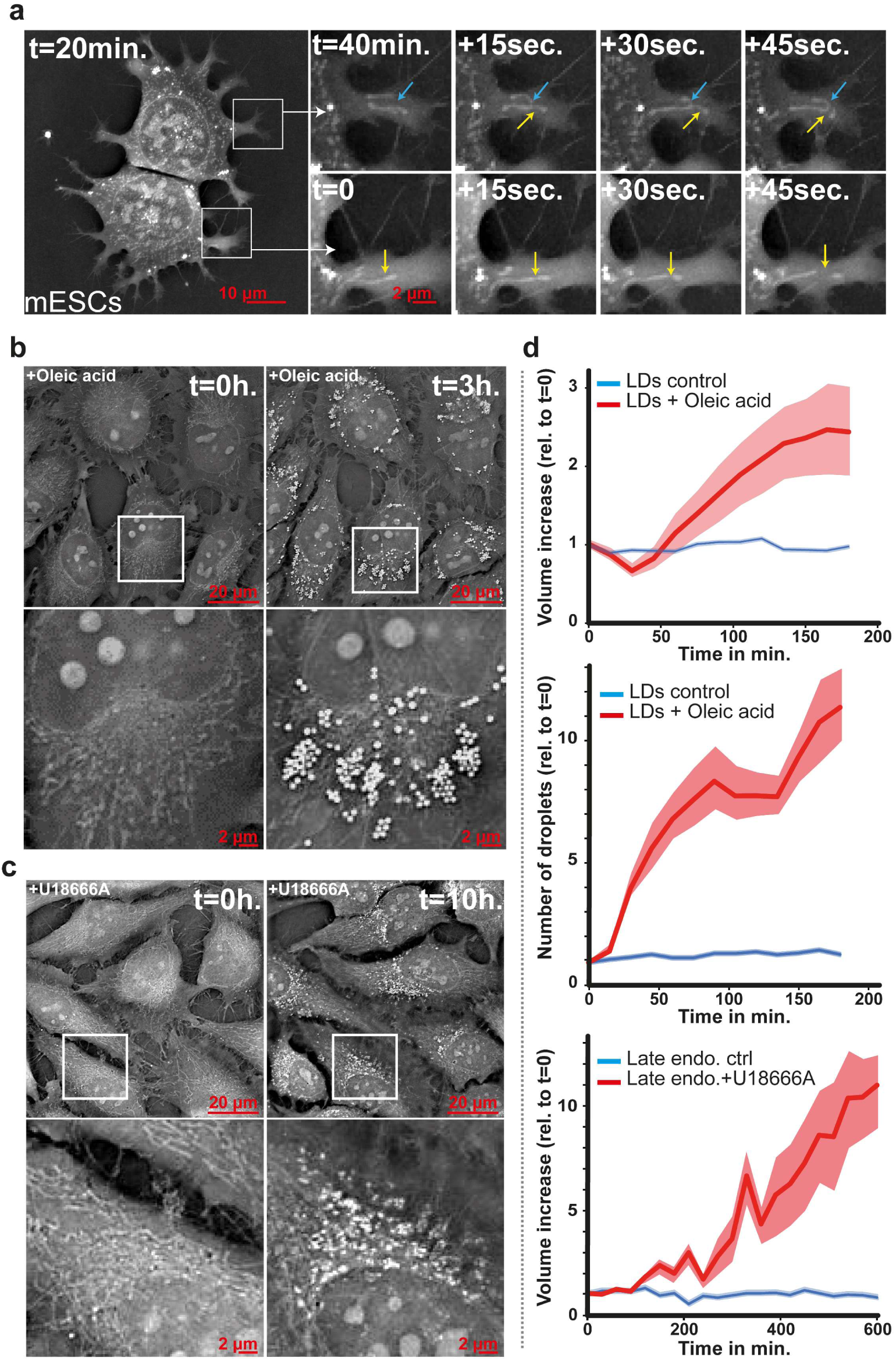
Holotomographic live microscopy reveals new lipid droplet and trafficking dynamics. **a**, Refractive index (RI) map of mouse embryonic stem cells times series showing mitochondrial fusion (blue arrow) and fission (yellow arrow) events **b**, Refractive index (RI) map of HeLa cells just before (t=0h) or three hours after (t=3h) loading with oleic acid (OA) or **c**, just before (t=0h) or ten hours after (t=10h) treatment with the compound U18666A. **d**, Quantification using computational image analysis of the increase of lipid droplet volume and number after OA loading, as well as of late endosomes accumulation after U18666A treatment (red curves, standard errors from multiple single lipid droplets or late endosomes shown in light red). All dynamics are compared to unperturbed conditions (blue curves, standard errors from multiple single lipid droplets or late endosomes shown in light blue).

Similarly, we followed lipid droplet (LD) dynamics overs hours using HTM at a frequency of 1 image.min^-1^. So far dynamics of LD have been visualized using fluorescence microscopy and Coherent Anti-Stokes Raman Scattering (CARS and the variant SRS)(15,16). Fluorescence microscopy coupled to specific bodipy dyes(15) provides contrast and specificity but is inherently phototoxic, limiting acquisition frequency and length to avoid significant perturbation of LD dynamics. CARS is a label-free technique; however, it is a complex and costly technology with limited signal specificity and temporal resolution(16). HeLa cells were incubated with oleic acid (OA) to trigger the growth of lipid droplets(17). The droplets could easily be segmented based on their RI values, which allowed to extract their features in 3D (S8 Fig). After three hours of incubation with OA, lipid droplets increased 8-fold in number and increased in size (Fig 3B). Droplets could be followed and quantified over time (Fig 3D and S4 movie). Because our HTM setup allows high frequency of acquisition and no phototoxicity, we could highlight unique dynamic features of cellular lipid accumulation. *i*.) The volume of lipid droplets significantly decreases in the first 30 minutes while the number of droplet increases suggesting that the cell prepares for lipid storage by first splitting existing droplets that can then grow. *ii*.) The average number of lipid droplets increases in a step wise manner: after 80 min, the number of LD reaches a plateau for app. 1h, during this time, the size of the LDs continues to increase. Then, the number of LDs increases strongly again, while the size of LDs progressively reaches a plateau. These dynamics suggest a feedback mechanism with size and number of lipid droplets as parameters, to reach an optimal intracellular lipid storage.

The fact that mitochondria, which are dense in membranes, and lipid droplets are readily observable by HTM, led us to speculate that lipid rich structures provide a clear RI signal. We therefore sought to induce the formation of a lipid rich organelle and we chose to do so by treating cells with U18666A, a charged amino derivative of cholesterol(18,19), a drug well-established to trigger lipid and cholesterol accumulation in the late endocytic pathway, and is used to mimic the phenotype of the Nieman Pick Type C disease. Using HTM at high frequency (1 image.min^-1^) relative to the acquisition length (10h), we indeed observed the appearance of perinuclear organelles with a well-defined RI signal. These structures are rich in cholesterol, as indicated by the filipin-positive staining, and co-localize with the late endosomal lipid LBPA (S9 Fig). The accumulation of these U18666A-induced structures was followed and quantified in 3D (Fig 3C, D and S5 video), showing that their volume increased app. 11-fold over a 10 h treatment. After a lag time of 100 minutes, U18666A-induced structure slowly appeared. At around 200 minutes, the increase in volume drastically increased barely reaching a plateau over the 10h period of the experiment. The absence of labeling, the spatial and temporal resolution of HTM should prove extremely useful to study the contribution of given genes in the formation of aberrant late endosomal structures in lysosomal storage diseases.

Finally, our unique capacity to observe multiple cellular organelles all at once within mouse embryonic stem cells (mESCs), and this over long time periods covering full cell cycles allowed us to capture a phenomenon of pre-mitotic organelle spinning. The rotation starts around 80 minutes before mitosis, occurs clock-wise at a mean speed of 5 degrees per minute (Fig 4 and S6-9 movie). To our knowledge, this is the first report of such phenomenon, which involves at least the nucleus(20), nucleoli, the nuclear membrane, lipid droplets and mitochondria (the global movement of the mitochondrial network is best seen in S6-9 movie) and suggests a potential mechanism of redistribution of the cellular material before division that needs to be investigated further.

**Fig 4.**
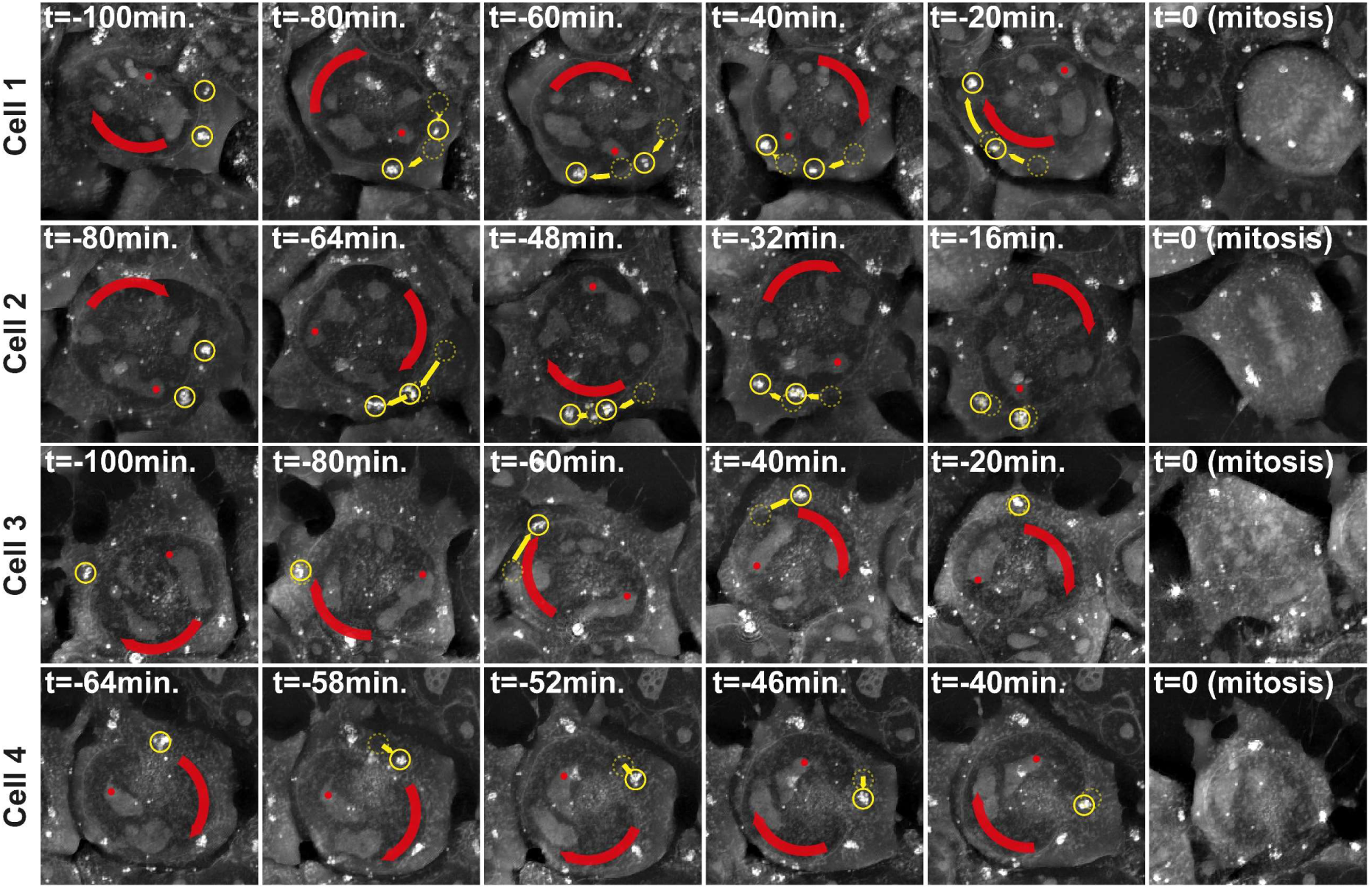
Holotomographic live microscopy reveals pre-mitotic organelle rotations in mouse embryonic stem cells. Time series of close-ups on mouse embryonic stem cells refractive index maps showing nuclear rotation (Red Arrow) and lipid droplet rotation (Yellow Arrow and circles) before mitosis. For a better feel of the rotation see Supplemental movies 6-9

This work demonstrates that the label-free nature and the low phototoxicity of HTM are advantageous to study the cell biology of specific organelles such as lipid droplets, mitochondria or pathologically lipid-loaded. We report to our knowledge for the first time a rotation of multiple organelles occurring in mESCs before mitosis, which could not be observed without the inherent multiplexing capacity of HTM (acquisition of multiple specific signals in one go). The potential role of such phenomenon in symmetric or asymmetric cellular division and cell fate needs to be unraveled. To do so using HTM to observe the effect of specific cell cycle signaling and cytoskeleton perturbations on the pre-mitotic organellar spinning seems to be a rational approach.

## Methods

### Cell cultures and seeding

Cells were cultured in MEM medium complemented with 10% FBS, 1% Pen/Strep, 1% L-glutamine and 1% non-essential amino acids. Cells were seeded for 24 h at low concentration on glass bottom Fluorodishes of 25mm and 0.17mm thickness (World Precision Instrument Inc.). For mouse embryonic stem cells, Fluorodishes were first coated with Vitronectin following the manufacturer’s protocol.

### Transfection

Cells were retrieved with trypsin from tissue culture dishes and seeded in Fluorodishes. After 24 h, the medium was changed, and the cells were transfected using Fugene (Promega) according to the manufacturer’s protocol.

### Cell fixation

Fixation was performed with paraformaldehyde (PFA) or cold methanol. For PFA fixation, cells were first washed 3x with PBS. Then, 2mL of PFA (3%) was added for 30 minutes at 37°C. Afterwards, dishes were washed 3x with PBS. Quenching of the preparations was performed with 50 mM NH4Cl in PBS at room temperature for 10 minutes before 3x PBS washes. Permeabilization was performed using 0.1% Triton X100 for 5 min at room temperature. In the case of methanol fixation, cells were washed 3x with PBS before adding pre-cooled methanol at −20°C for 4 min. Cells were washed 3x with PBS after fixation.

### Immunofluorescence

Cells were seeded in Fluorodishes for 48 h (if so, including transfection) prior to fixation and permeabilization. Overnight blocking was performed in PBS with 0.5% BSA. Primary and secondary antibodies were applied for 30 minutes at room temperature each with in between 3x washes of PBS-0.5% BSA for 5 min. Finally, the preparation was washed again three times with PBS-0.5% BSA and post-fixed for 15 min at room temperature with 3% PFA followed by 3x PBS washes. Hoechst (Invitrogen) was used at 2µg/mL for 30 minutes at room temperature, BODIPY (Invitrogen) at 1µg/mL for 30 minutes in physiological conditions. Filipin (Sigma) was used at a dilution of 1:50 (from stock 50µg/ml).

### Drug treatments

U18666A was applied to the cells at a dilution of 1:2000 (from a stock of 10 mg/ml). Oleic acid was applied at a dilution of 1:5000 (from stock 1mg/mL).

### Imaging

Holo-tomographic Microscopy (HTM) in correlation with epifluorescence was performed on the 3D Cell-Explorer Fluo (Nanolive) using a 60x air objective (NA = 0.8) at a wavelength of λ=520nm (Class1 low power laser, sample exposure 0.2 mW/mm^2^) and USB 3.0 CMOS Sony IMX174 sensor / Quantum E ciency (typical) 70 % (at 545 nm) / Dark Noise (typical) 6,6 e- / Dynamic Range (typical) 73,7 dB. The correlative acquisitions with bright-field, phase contrast and DIC were done on an Axiovert 200M (Zeiss) using a 63x objective (NA 1.4).

### Live cell imaging

Physiological conditions for live cell imaging were reached with a top-stage incubator from oko-lab. This solution is compatible with the microscope stage and allows for a complete control of the temperature and percentage of CO2 inside the chamber and thus for the cells. A constant temperature of 37°C, an air humidity saturation as well as a level of 5% CO2 was achieved throughout the acquisitions.

### Lipid droplets and late endosomes analysis

Cells are imaged by HTM as described above. From the HTM acquisition, a digital segmentation of the cell surface layer and of the lipids is performed based on the refractive index values. This digital stain is exported from the HTM microscope software in a .tiff format before being imported into ImageJ. There, a plugin was coded for lipid droplets analysis (https://github.com/MatAtNanolive/SimpleFIJImacro-). The algorithm relies on two existing available plugins for ImageJ. The first, Object Counter3D(21), performs a particle analysis, while the second, MorphoLibJ(22) offers a watershed segmentation. The whole method, as described below, is applied to the imported images. First, it separates the two channels from the segmentation. Then, Object Counter3D performs a particle analysis in the channel of the segmented lipid droplets (now seen as particles thanks to the digital stain). It extracts multiple parameters (such as coordinates in x,y,z and volume). Afterwards, the watershed segmentation is run with the MorphoLibJ plugin based on the channel of the cell surface segmentation. This step is semi-automatic as the user draws points with a 4-pixels diameter inside the cells and with a 20-pixels diameter in the background which guides the segmentation and enables to make match lipid droplets with their respective cell. Measurements and data extraction are done automatically by the plugin and saved into excel sheets containing all the information about the individual lipid droplets and the distribution of the lipids into the cell.

## Acknowledgement

We thank the LipidX consortium and the NCCR chemical biology for funding this work. We thank José Artacho from EPFL’s Imaging Platform for his great help with the phase contrast and DIC correlative acquisitions. In addition, we thank Hubert Becker for hand-labelling of mitochondria and lipid droplets, Aleksandra Mandic for providing mESC, Kristina Shoonjans for the Hepa 1.6 cells and Pierre Gönczy for the U2OS cells, Sebastian Jessberger for KDEL-GFP, Anne-Laure Mahul for Mito-YFP, Tom Kirchhausen for Emerin-GFP and Jean Grunberg for anti-LBPA.

## Data Availability

The data supporting this study are available from the corresponding authors upon reasonable request.

---

**S1 Fig. Comparison of holotomographic microscopy and other label-free microscopy techniques.**

**S2 Fig. Impact of methanol or paraformaldehyde fixation of HeLa cells on holotomographic microscopy results.**

**S3 Fig. Aspect of various cell lines when observed with holotomographic microscopy.**

**S4 Fig. Comparison of specific fluorescent signals and refractive index map**. Visual comparison of a, HeLa cells refractive index (RI) map to Golgi apparatus fluorescent signal (NAGTI-GFP) or b, to an endoplasmic reticulum fluorescent signal (KDEL-GFP) or c, to a late endosome accumulation signal (Filipin).

**S5 Fig. Correlation of various fluorescent and RI signal using various metrics. a**, Bootstrapped Kolmogorov-Smirnov test of the distribution of refractive index (RI) values under specific fluorescent signals against the global cellular RI distribution. Statistical test p-values are plotted as a function of the stringency of the threshold used to define the fluorescence mask, from a specific or random fluorescent object signal. b, Pearson correlation of various fluorescent and RI signals.

**S6 Fig. Full width half maximum quantification of minimal mitochondrial thickness using FIJI software**. The mitochondrial thickness has been measured within fibroblasts as the width observed at the half maximum of the mitochondrial signal distribution. The signal distributions along transversal lines have been defined within the FIJI software on zero padded image and are represented in the red enlargement squares (red lines 1-3).

**S7 Fig. Comparison of refractive index maps of cells when transfected with fluorescently-tagged mitochondrial protein**. Refractive index (RI) map of HeLa cells a, after transfection with a neutral mitochondrial Ds-Red marker or b, with the mitochondrial fusion protein Fis1-GFP.

**S8 Fig. Object segmentation within refractive index 3D map using FIJI. a**, refractive index map of HeLa cells. b, Lipid droplet segmentation using FIJI displayed in 2D on the central layer of the 3D sample volume. Scale bar, 20 µm.

**S9 Fig. Comparison between refractive index map and late endosome or cholesterol fluorescent signal**. Compared to control, cells treated with U18666A show different perinuclear structures in the refractive index (RI) map that overlap with LBPA immunostaining and cholesterol filipin staining, indicating that the accumulation of cholesterol-rich late endosomes after treatment is observable in the RI map.

**S1 movie. Live imaging of mitochondrial fission and fusion events in mouse embryonic stem cells using holotomographic microscopy**. movie length, 6h, acquisition frequency, 1 image / 15 seconds. Related to figure 3a.

**S2 movie. Live imaging of mitochondrial fission in HeLa cells using holotomographic microscopy**. Fission events are shown by red arrows, movie length, 1h, acquisition frequency, 1 image / 30 seconds. Related to figure 3a.

**S3 movie. Live imaging of mitochondrial fusion in HeLa cells using holotomographic microscopy**. Fusion events are shown by blue arrows, movie length, 1h, acquisition frequency, 1 image / 30 seconds. Related to figure 3a.

S4 movie. Live imaging of lipid droplet growth in HeLa cells after oleic acid loading using holotomographic microscopy. movie length, 3h, acquisition frequency, 1 image.min^-1^. Related to figure 3b.

**S5 movie. Live imaging of late endosome accumulation in HeLa cells after U18666A treatment using holotomographic microscopy**. movie length, 10h, acquisition frequency, 1 image.min^-1^. Related to figure 3c.

**S6 movie. Live imaging of pre-mitotic nuclear rotation in mouse embryonic stem cells using holotomographic microscopy**. movie length, 5h30, acquisition frequency, 1 image.min^-1^. Related to figure 4.

**S7 movie. | Live imaging of pre-mitotic nuclear rotation in mouse embryonic stem cells using holotomographic microscopy**. movie length, 5h30, acquisition frequency, 1 image.min^-1^. Related to figure 4.

**S8 movie. Live imaging of pre-mitotic nuclear rotation in mouse embryonic stem cells using holotomographic microscopy**. movie length, 1h45, acquisition frequency, 1 image.min^-1^. Related to figure 4.

**S9 movie. Live imaging of pre-mitotic nuclear rotation in mouse embryonic stem cells using holotomographic microscopy**. movie length, 1h45, acquisition frequency, 1 image.min^-1^. Related to figure 4.

